# First detection of tick-borne encephalitis virus in *Ixodes ricinus* ticks and their rodent hosts in Moscow, Russia

**DOI:** 10.1101/480475

**Authors:** Marat Makenov, Lyudmila Karan, Natalia Shashina, Marina Akhmetshina, Olga Zhurenkova, Ivan Kholodilov, Galina Karganova, Nina Smirnova, Yana Grigoreva, Yanina Yankovskaya, Marina Fyodorova

**Affiliations:** Central Research Institute of Epidemiology, Novogireevskaya st 3-A, 415, Moscow, Russia, 111123; Scientific Research Disinfectology Institute, Nauchniy proezd st. 18, Moscow, Russia, 117246; Chumakov Institute of Poliomyelitis and Viral Encephalitides (FSBSI “Chumakov FSC R&D IBP RAS), prem. 8, k.17, pos. Institut Poliomyelita, poselenie Moskovskiy, Moscow, Russia, 108819; Institute for Translational Medicine and Biotechnology, Sechenov University, Bolshaya Pirogovskaya st, 2, page 4, room 106, Moscow, Russia, 119991; Leninskie Gory st. 1-12, MSU, Faculty of Biology, Moscow, Russia, 119991; Pirogov Russian National Research Medical University, Ostrovityanova st. 1, Moscow, Russia, 117997

**Keywords:** Tick-borne encephalitis virus, *Ixodes ricinus*, *Borrelia burgdorferi*, Vector-borne diseases, Small mammals

## Abstract

Here, we report the first confirmed autochthonous tick-borne encephalitis case diagnosed in Moscow in 2016 and describe the detection of Tick-borne encephalitis virus (TBEV) in ticks and small mammals in a Moscow park.

The paper includes data from two patients who were bitten by TBEV-infected ticks within the Moscow city limits; one of these cases led to the development of the meningeal form of TBE. Both TBEV-infected ticks attacked patients in the same area. We collected ticks and trapped small mammals in this area in 2017. All samples were screened for the presence of pathogens causing tick-borne diseases by PCR. The TBEV-positive ticks and small mammals’ tissue samples were subjected to virus isolation. The sequencing of the complete polyprotein gene of the positive samples was performed.

A total of 227 questing ticks were collected. TBEV was detected in five specimens of *Ixodes ricinus*. We trapped 44 small mammals, mainly bank voles (*Myodes glareolus*) and pygmy field mice (*Apodemus uralensis*). Two samples of brain tissue from bank voles yielded a positive signal in RT-PCR for TBEV. We obtained six virus isolates from the ticks and brain tissue of a bank vole. Complete genome sequencing showed that the obtained isolates belong to the European subtype and have low diversity with sequence identities as high as 99.9%. GPS tracking showed that the maximum distance between the exact locations where the TBEV-positive ticks were collected was 185 m. We assume that the forest park was free of TBEV and that the virus was recently introduced.

## Introduction

Tick-borne encephalitis virus (TBEV) is a flavivirus that is transmitted by the bite of infected hard-bodied ticks. The geographic range of TBEV is mainly in Eurasia spanning Japan to the Russian far east towards Central Europe, corresponding to the distribution of the ixodid ticks (Lindquist and Vapalahti, 2008; Süss, 2011; Zlobin, 2010). The prevalence of the virus, as well as its distribution, can increase based on the different yearly factors governing the different parts of the vectors’ and hosts’ ranges (Korenberg, 2009). In particular, changes in the boundaries of the TBE risk areas were described in Germany (Süss et al., 2004) and Russia (Tokarevich et al., 2011). Furthermore, the detection of new TBEV foci has been reported in Sweden (Haglund, 2002; Jaenson et al., 2012) and Norway (Csango et al., 2004; Haglund, 2002; Skarpaas et al., 2006, 2004). There is evidence regarding the reactivation of old foci of TBEV, which for a long time (decades) have been inactive, thus turning the places into a risk area (Frimmel et al., 2014; Jääskeläinen et al., 2006; Kristiansen, 2002).

In Russia, out of 85 regions, 47 belong to TBE risk areas (“On the list of endemic territories for viral tick-borne encephalitis in 2015,” 2015), but Moscow and most districts of the Moscow Oblast are not endemic for TBEV. Only two districts in the north of the Moscow Oblast are included in the TBE risk area (“On the list of endemic territories for viral tick-borne encephalitis in 2015,” 2015; Shevtsova et al., 2008), but local cases of the disease have not been recorded there for several decades. The Moscow Oblast borders two TBE endemic regions in the north and northwest with the Tver Oblast and Yaroslavl Oblast, respectively (Fig. 1). The most recent, confirmed cases of TBE in the Moscow Oblast are from 1948-1954. At the time of the outbreak, at least 150 patients were documented in two regions lying to the northwest of the Moscow Oblast (Drozdov, 1956). In the city of Moscow, local cases of the disease were not registered. The city has experienced a significant population increase in the past, growing due to the expansion of the surrounding territories. Some of the new urban areas were not developed, and they were preserved as forest parks. The local fauna of such forest parks has remained and includes small mammals and ticks (Uspensky, 2017). Surveys of the forest parks of Moscow showed the presence of *Ixodes ricinus* in the majority of the city’s parks (Shashina and Germant, 2015). A recent study of the ticks collected in Moscow’s parks revealed the prevalence of *Borrelia burgdorferi* s.l., *Anaplasma phagocytophilum* and *Ehrlichia sp*., but TBEV was not detected (Shashina and Germant, 2015). In Moscow, 5-14 clinical TBE cases have been recorded annually, but none of them have been autochthonous (*On the incidence of tick-borne borreliosis in Moscow in 2014*, 2015; Yankovskaya et al., 2017).

**Fig. 1.**
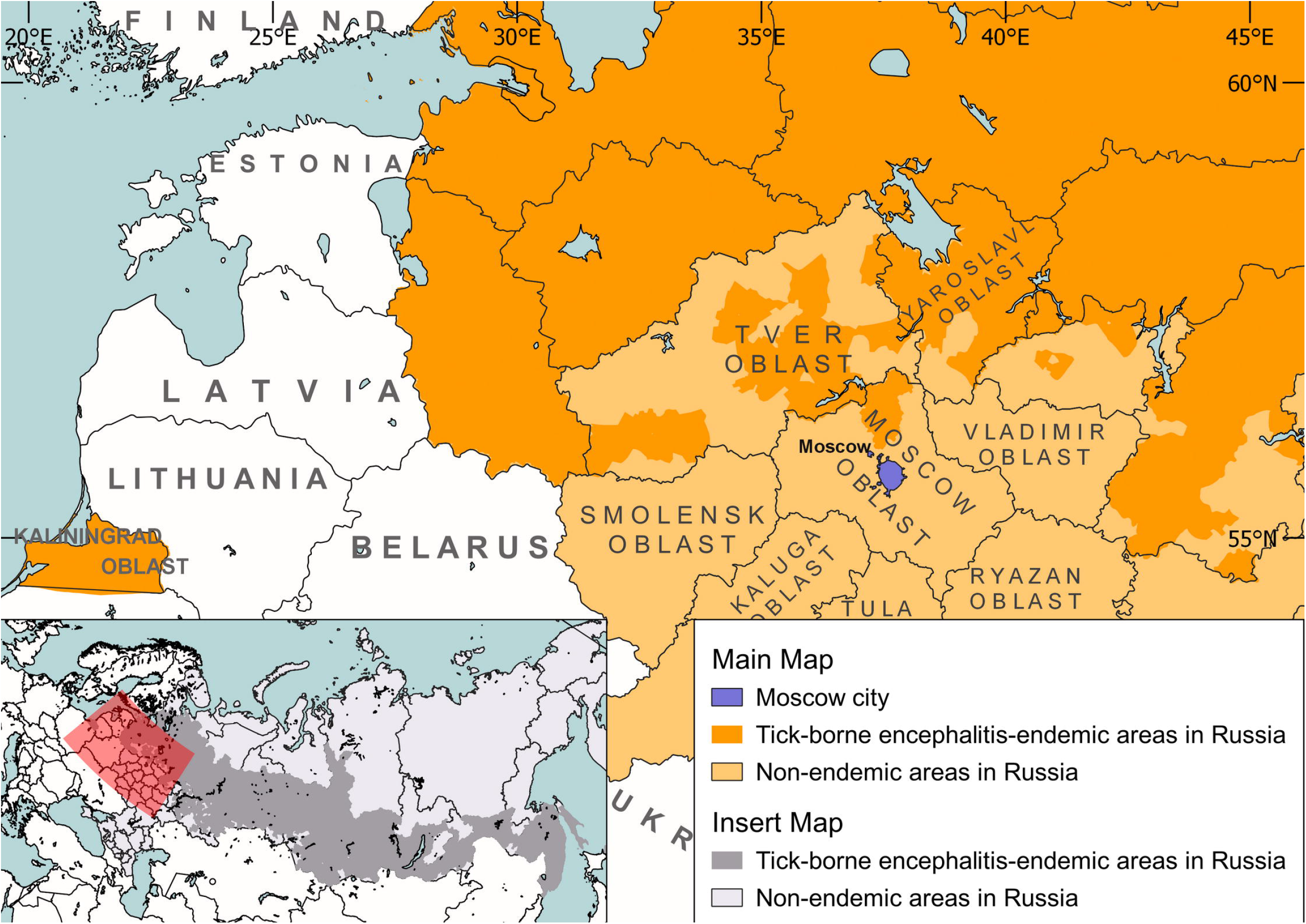
Tick-borne encephalitis-endemic areas in Russia (Based on (*On the list ofendemic territories for viral tick-borne encephalitis in 2015*, 2015)).

Therefore, we aimed to report the confirmed autochthonous TBE case diagnosed in Moscow in 2016 and to study the prevalence of TBEV among ixodid ticks and small mammals in an urban forest park. In addition, this paper seeks to identify the origin of the discovered TBE foci and to determine whether the described cases are due to the reactivation of formerly active foci or if there is evidence of introduction of TBEV from a new territory.

## Methods

### Patients

The paper includes data from two patients who were bitten by TBEV-infected ticks within the Moscow city limits. We interviewed both patients and considered their medical records. For the patient described in Case-2, blood samples were collected on the 17th and 41st days after the tick bite to test for anti-TBEV IgM and IgG.

### Research area

We obtained first two TBEV-infected ticks from the same area in the western part of the city. Hence, we focused on forest parks from this area. These parks adjoin the forests of the Moscow Oblast and the Moscow Ring Road (a large highway with busy round-the-clock traffic) separates them. We divided the research area into three study sites (Fig. 2). Site-1 was the main site used for field research since in this forest park, a TBEV-infected tick attacked a patient in July 2017 (see Case-2 below). Additionally, we selected Site-2 and Site-3 north and west of Site-1, respectively, to check the areas in the city boundaries. The forest parks on Site-1, Site-2, and Site-3 are free of regulation and management services (such as grass mowing, deratization, spraying acaricides, etc.). The areas of these sites are 104 ha, 218 ha and 179 ha, respectively. The local people actively use these parks for walking, physical activity, recreation, and dog walking.

**Fig. 2.**
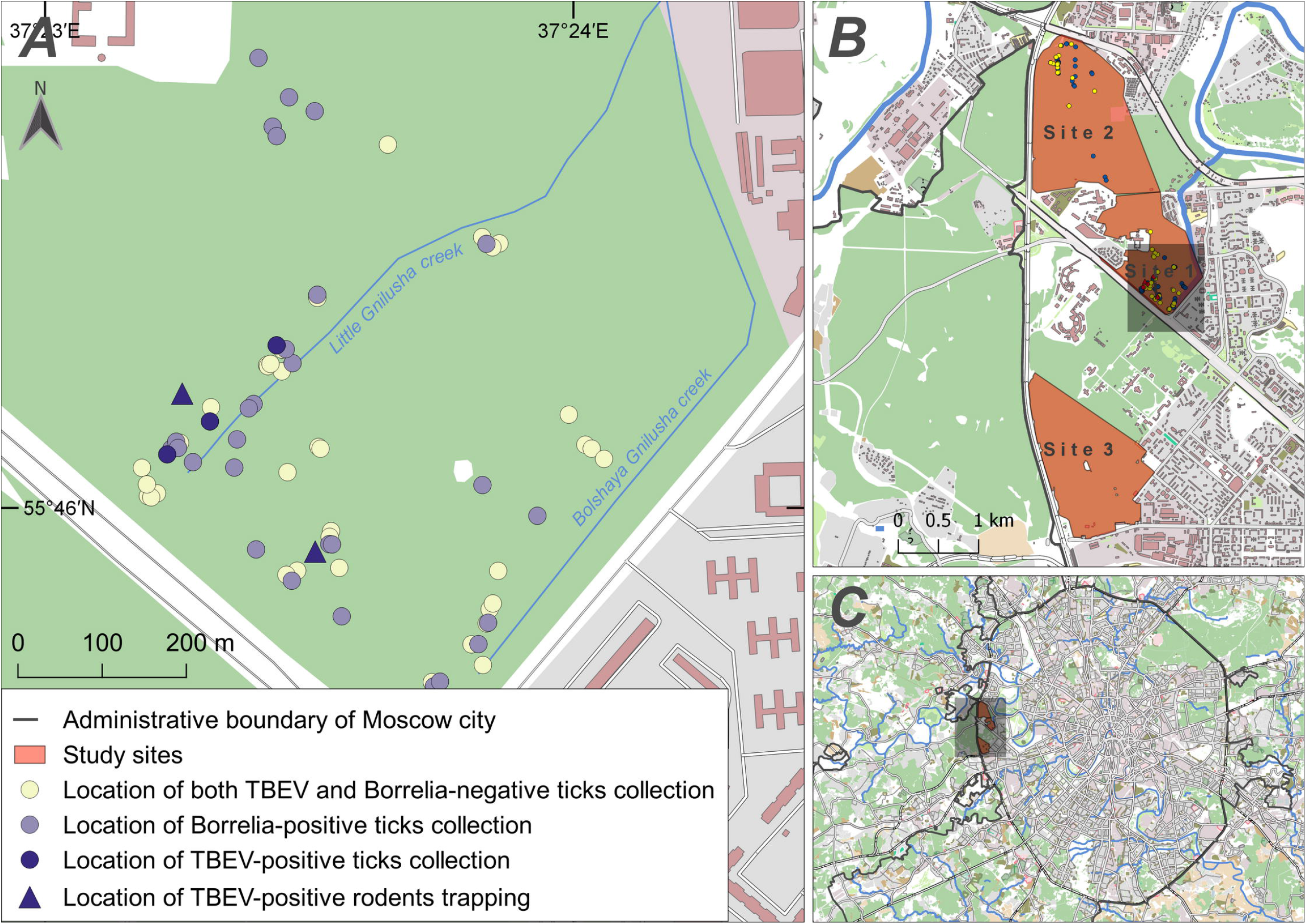
Study area with depicted locations of tick collection and small mammal trapping (A). The inserted maps represent the location of the study sites (B) and the administrative boundary of Moscow city (C).

### Tick collection

Ticks were collected through flagging on regular transects in May 2017 (one sample session) and in August-September 2017 (2-3 times per week). Any tick collected was placed in a separate tube. We recorded the GPS coordinates of each tick collection site. All tick specimens were identified to stage and species by morphological characteristics (Filippova, 1977). Additionally, we performed molecular species identification (see below for details).

### Small mammals

Small mammals were captured in October 2017 using live traps (Shchipanov, 1987) baited with sunflower oil and oat. The traps were set up for four nights placed at 5 m intervals along two linear transects. The total capture effort was 100 traps per nights. We chose the location for small mammals trapping based on the GPS tracking of the TBEV-positive tick collection sites.

### Extraction, Molecular detection and genotyping of pathogens

All the collected ticks were tested for the presence of tick-borne pathogens by a real-time PCR assay. Initially, the ticks were washed consistently in 70% alcohol and 0.15 M NaCl solution, and then individual ticks were homogenized with TissueLyser LT (Qiagen, Germany). Thereafter, DNA/RNA was extracted from 100 μl of the tick suspension using a commercial kit AmpliSens RIBO-prep kit (Central Research Institute of Epidemiology, Moscow, Russia) following the manufacturer’s instructions. PCR-based detection of TBEV, *Borrelia burgdorferi* sl, *Anaplasma phagocytophilum, Ehrlichia chaffeensis*, and *Ehrlichia muris* was performed using a commercial multiplex PCR kit (AmpliSens TBEV, *B. burgdorferi* s.l., *A. phagocytophilum, E. chaffeensis/E. muris*-FL; Central Research Institute of Epidemiology, Moscow, Russia) according to the manufacturer’s instructions. Each run included a negative control and the mix of the five positive recombinant DNA controls of the TBEV 5’UTR-C gene fragment; *B. burgdorferi* sl and *E. chaffeensis, E. muris* 16S gene fragments, *A. phagocytophilum* msp2 gene fragment and RNA internal control with 10^5^ copies/ml. The RNA template was reverse transcribed using Reverse Transcriptase (Thermo Scientific). The commercial PCR kit AmpliSens *B. miyamotoi*-FL (Central Institute of Epidemiology, Moscow, Russia) was used according to the manufacturer’s instructions for the *Borrelia miyamotoi* glpQ gene.

To check the species diagnoses based on the morphological characteristics, we performed qPCR (Table 1). Furthermore, for tick identification, we sequenced the *ITS-2* gene fragments and *COI* gene fragments (Table 1). PCR products of the appropriate length were separated by electrophoresis on a 1.5% agarose gel supplemented with ethidium bromide. Amplicon purification was performed using the RIBO-prep extraction kit (AmpliSens, Russia).

**Table 1.**
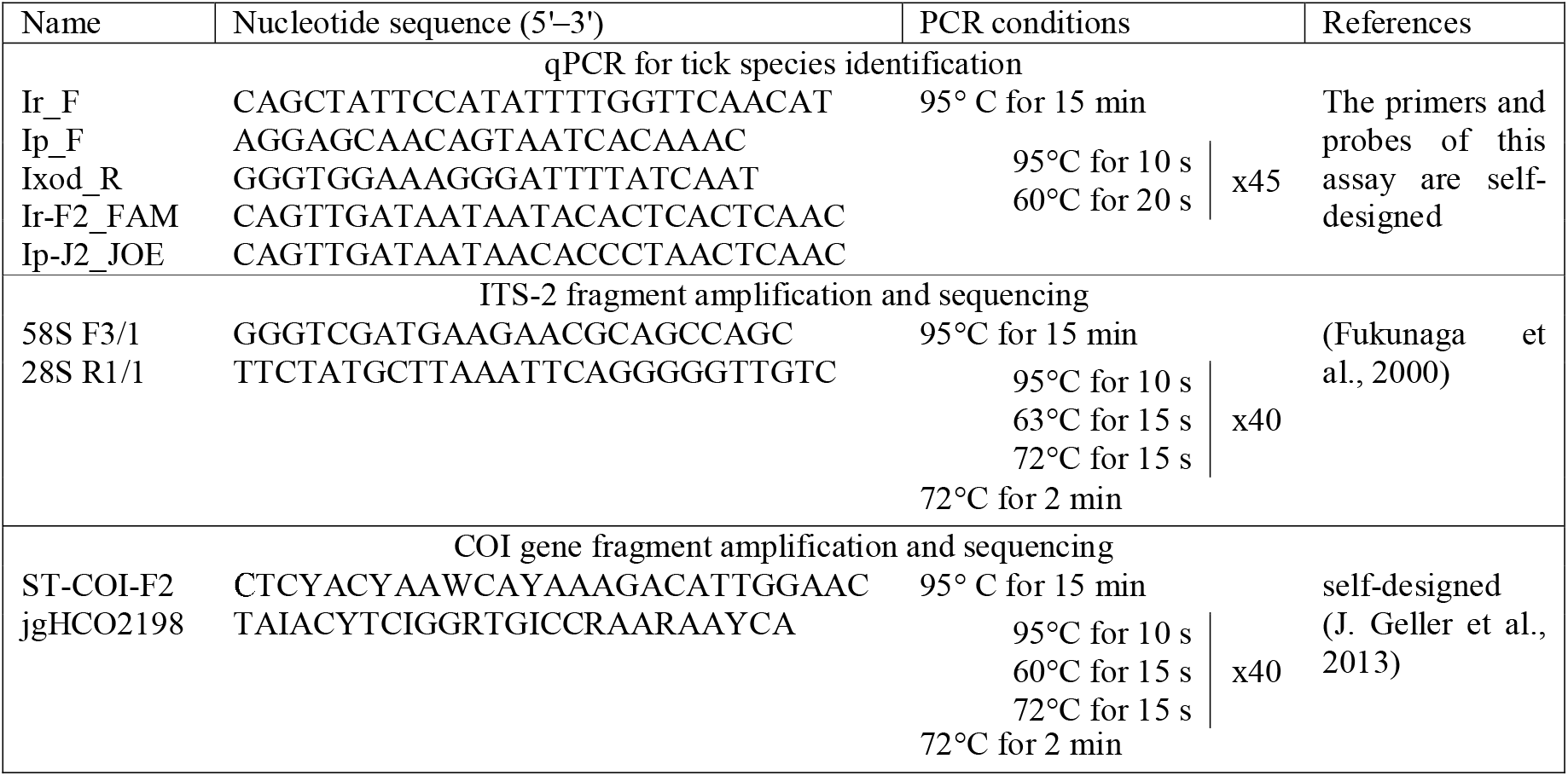
Oligonucleotide sequences of the primers and probes used in PCR amplification and sequencing

### Virus isolation

The TBEV-positive ticks and small mammals’ tissue samples were subjected to virus isolation. For this, we used the pig embryo kidney (PEK) cell line and suckling mice. The PEK cell line was maintained at 37°C in medium 199 (PIPVE, Russia) supplemented with 5% FBS (Gibco). The suckling mice (FSBI, Scientific Center of Biomedical Technologies, Stolbovaya Branch, Russia) were infected intracerebrally with the supernatant obtained from the PCR-positive tick and tissue sample suspensions. The mice were sacrificed at the appearance of signs of the disease or on the 14th day of the experiment. The mice were maintained according to the international guidelines for animal husbandry, including the recommendations of CIOMS, 1985 and the FELASA Working Group Report, 1996–1997.

### 50% plaque reduction neutralization test (PRNT50)

We tested rodents’ serum with neutralizing antibody for TBEV using a 50% plaque reduction neutralization test (PRNT50). The analysis was performed as described earlier (Pripuzova et al., 2013). For this test, we used strain MOS-152-T-2017, isolated from the tick that bit the patient described in Case-2. Briefly, sequential dilutions of the sera samples were prepared in 199 medium on Earle solution with the addition of 2% FBS (Gibco). An equal volume of viral suspension containing 40-70 plaque forming units was added to the sera dilution. Virus-positive sera were incubated at 37°C for 1 h, added to PEK cell monolayers, and incubated for one hour for virus adsorption. Then, each well was overlaid with 1% Bacto-agar (Difco) on Earle solution containing 7.5% FBS and 0.015% neutral red. Every experiment included appropriate controls, i.e., negative and positive murine sera with known titers. The antibody titer was calculated according to the modified Reed and Muench method (Reed and Muench, 1938).

### The sequencing

We used RNA isolated from tick suspensions and rodents’ brain tissue for sequencing. The purified PCR products were sequenced bidirectionally on an ABI Prism 3500 (Applied Biosystems, USA). The sequences obtained were deposited in NCBI GenBank under the following accession numbers: TBEV MH663426-MH663428 (complete polyprotein gene, 9521 bp) and MH663429-MH663430 (NS1 protein gene, 1054 bp). The partial sequence alignment and phylogenetic tree construction were performed using the software MEGA7.

### Data analysis

We conducted the data analysis in R using the package PropCIs (Scherer, 2014). The sites where the questing ticks were collected and the small mammals were trapped were mapped using Quantum GIS free software. The nucleotide sequences were aligned, compared, and analyzed using MEGA7, ClustalW, and BLAST.

## Results

### Patient cases

**Case 1.** A 43-year-old man was bitten by a tick in a forest park in the west of Moscow on August 8, 2016. The patient stated that he had not had any other tick bites and had not visited TBE-endemic areas. The patient also declined TBE vaccination. On August 13, fever, headache, and myalgia developed in the patient. Erythema migrans were absent at the site of the tick bite. The patient started antibiotic treatment (doxycycline) on an outpatient basis after consulting a doctor. On August 29, the patient was hospitalized with high fever (temperature 40.0°C). ELISA tests on anti-TBE IgM and IgG showed positive results. The diagnosis was Tick-borne encephalitis, meningeal form.

**Case 2.** On July 25, 2017, a 27-year-old woman reported a tick bite after walking in a forest park in the west of Moscow. The patient brought the tick to a commercial laboratory to test. RT-PCR yielded a positive result for TBEV. On July 27, the patient received prophylactic immunoglobulin therapy (1:80, 6 ml). Two ELISAs of serum samples for anti-TBE IgM and IgG conducted on the 17^th^ and 41^st^ days after the tick bite showed negative results. No symptoms of acute infection manifested.

### Questing ticks

The questing ticks field assays resulted in the collection of 227 ticks from vegetation in the study area, and 225 specimens out of the total collection were identified morphologically and through qPCR to be *Ixodes ricinus* (Table 2). Only one tick was identified as *I. persulcatus* (female), and one was identified as *Dermacentor reticulatus* (male). The mean abundance of adult *I. ricinus* at Site-1 varied in August from 1,6 to 3,0 ticks/km (Table 3). We did not estimate the abundance of ticks at Site-2 and Site-3.

**Table 2.** Questing ixodid ticks collected in the study area

| Location | *I. ricinus* |  | *I. persulcatus* | *D. reticulatus* | Total |
| --- | --- | --- | --- | --- | --- |
|  | Adults | Nymphs |  |  |  |
| Site-1 | 106 | 42 | 1 | 0 | 149 |
| Site-2 | 47 | 10 | 0 | 1 | 58 |
| Site-3 | 17 | 3 | 0 | 0 | 20 |

**Table 3.**
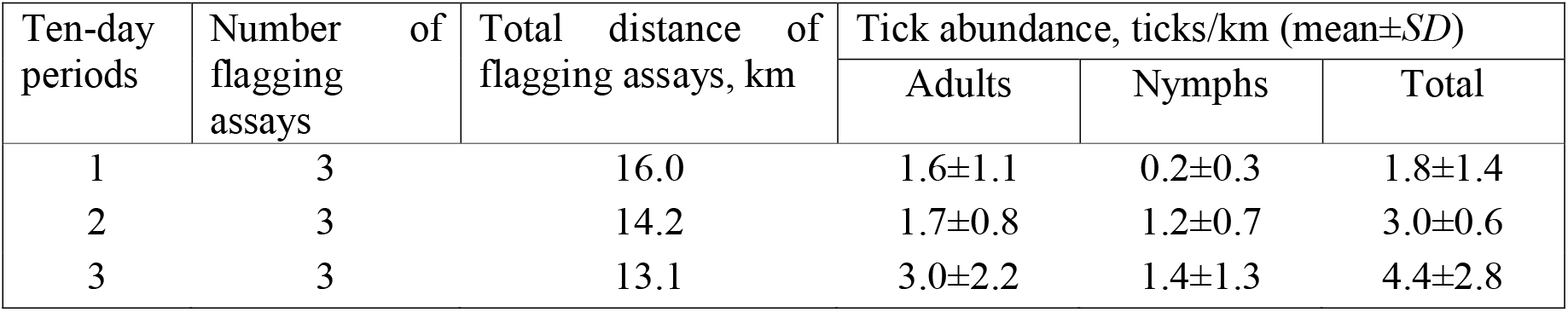
The abundance of *I. ricinus* at Site-1, August 2017

TBEV was detected in five specimens, including two adult males, two adult females, and one nymph of *I. ricinus* (Table 4). All TBEV-positive ticks were collected at Site-1. We determined the exact coordinates of the collection site for the three TBEV-positive ticks. Mapping showed that all the TBEV-positive ticks were obtained at the same location along the small river on the western part of the park (Fig. 2). The maximum distance between the TBEV-positive ticks was 185 m. The mean prevalence of *Borrelia burgdorferi* s.l. in ticks at all the studied sites was 32,4%, the difference between Site-1 and Site-2 was not significant (*χ^2^* = 1.58, *df* = 1, *p* = 0.21).

**Table 4.** Prevalence of TBEV and *B. burgdorferi sl* in collected *Ixodes ricinus* ticks

| Stage | Total | Number of PCR-positive samples, Frequency (% – CI 95%) |  |
| --- | --- | --- | --- |
|  |  | TBEV | *B. burgdorferi* s.l. |
| Site-1 | 149 | 5 (3.4% – 1.1–7.7%) | 54 (36.2% – 28.5–44.5%) |
| Adults: | 107 | 4 (3.7% – 1.0–9.3%) | 42 (39.3% – 30.0–49.2%) |
| Female | 52 | 2 (3.8% – 0.5–13.2%) | 22 (42.3% – 28.8–56.8%) |
| Male | 55 | 2 (3.6% – 0.4–12.5%) | 20 (36.4% – 23.8–50.4%) |
| Nymph | 42 | 1 (2.4% – 0.1–12.6%) | 12 (28.6% – 15.7–44.6%) |
| Site-2 | 58 | 0 (0.0%) | 15 (25.9% – 15.3–39.0%) |
| Adults: | 48 | 0 (0.0%) | 15 (31.3% – 18.7–46.3%) |
| Female | 23 | 0 (0.0%) | 7 (30.4% – 13.2–52.9%) |
| Male | 25 | 0 (0.0%) | 8 (32.0% – 14.9–53.5%) |
| Nymph | 10 | 0 (0.0%) | 0 (0.0%) |
| Site-3 | 20 | 0 (0.0%) | 7 (35.0% – 15.4–59.2%) |
| Adults: | 17 | 0 (0.0%) | 4 (23.5% – 6.8–49.9%) |
| Female | 14 | 0 (0.0%) | 3 (21.4% – 4.7–50.8%) |
| Male | 3 | 0 (0.0%) | 1 (33.3% – 0.8–90.6%) |
| Nymph | 3 | 0 (0.0%) | 3 (100.0%) |

### Small mammals

Out of the 44 rodents that were live-trapped, 13 of these were released since they were not fully mature (individuals without previous breeding experience) and had no ticks on them (Table 5.). At Site-1, we trapped small mammals of four species; the sample included mainly bank voles (*Myodes glareolus*) and pygmy field mice (*Apodemus uralensis*). From the trapped animals, we collected larvae of *I. ricinus* and larvae and adult females of *Ixodes trianguliceps* (Table 5). In addition to the ixodid ticks, we found a large number of chiggers identified as *Hirsutella zachvatkini*.

**Table 5.** The number of live-trapped small mammals and the number of ixodid ticks collected from the investigated animals

|  | Total | *Myodes glareolus* | *Apodemus uralensis* | *Apodemus agrarius* | *Sorex aureus* |
| --- | --- | --- | --- | --- | --- |
| Small mammals |  |  |  |  |  |
| Trapped | 44 | 20 | 21 | 1 | 2 |
| Collected for further work | 30 | 20 | 9 | 1 | 0 |
| Prevalence of ticks collected from small mammals, the number and % (CI95%) |  |  |  |  |  |
| *I. ricinus* larvae | 3<br>9.7%<br>(2.0–25.8%) | 1<br>5.3%<br>(0.1–26.0%) | 1<br>11.1%<br>(0.3–48.2%) | 1<br>100.0% | – |
| *I. trianguliceps* larvae | 11<br>35.5%<br>(19.2–54.6%) | 3<br>15.8%<br>(3.4–39.6%) | 8<br>88.9%<br>(51.8–99.7%) | 0<br>0.0% | – |
| *I. trianguliceps* female | 1<br>3.2%<br>(0.1–16.7%) | 0<br>0.0% | 1<br>11.9%<br>(0.3–48.2%) | 0<br>0.0% | – |

We tested samples of small mammalian tissues for the presence of TBEV RNA and DNA of *B. burgdorferi* s.l., *B. miyamotoi, Leptospira sp., Bartonella sp., Rickettsia sp., Coxiella burnetti, A. phagocytophilum, E. chaffeensis*, and *E. muris*. Two specimens of brain tissue of bank vole (*M. glareolus*) yielded a positive signal in RT-PCR for TBEV. Furthermore, we found DNA of *Borrelia burgdorferi* s.l., *Leptospira sp., Bartonella sp*., and *Rickettsia sp*. (Table 6). PRNT50 showed that 24 of the 28 studied rodents had neutralizing antibodies for TBEV (Table 7). The distance between TBEV-positive rodent trapping locations was 245 m (Fig. 2).

**Table 6.** Prevalence of vector-borne and zoonotic pathogens in the tissue samples of the investigated small mammals

| Pathogens | Number of PCR-positive samples, Frequency (% – CI 95%) |  |
| --- | --- | --- |
|  | *Myodes glareolus* | *Apodemus uralensis* |
| The total number of tested animals | 20 | 9 |
| TBEV | 2 (10.0% – 1.2–31.7%) | 0 |
| *Borrelia burgdorferi* s.l. | 8 (40.0% – 19.1–63.9%) | 7 (77.8% – 40.0–97.2%) |
| *Borrelia miyamotoi* | 0 | 0 |
| *Bartonella* spp. | 7 (35.0% – 15.4–59.2%) | 4 (44.4% – 13.7–78.8%) |
| *Leptospira* spp. | 0 | 1 (11.1% – 19.1–63.9%) |
| *Rickettsia* spp. | 0 | 1 (11.1% – 19.1–63.9%) |
| *Coxiella burnetii* | 0 | 0 |
| *Anaplasma phagocytophilum* | 0 | 0 |
| *Ehrlichia chaffeensis* / *E. muris* | 0 | 0 |

**Table 7.**
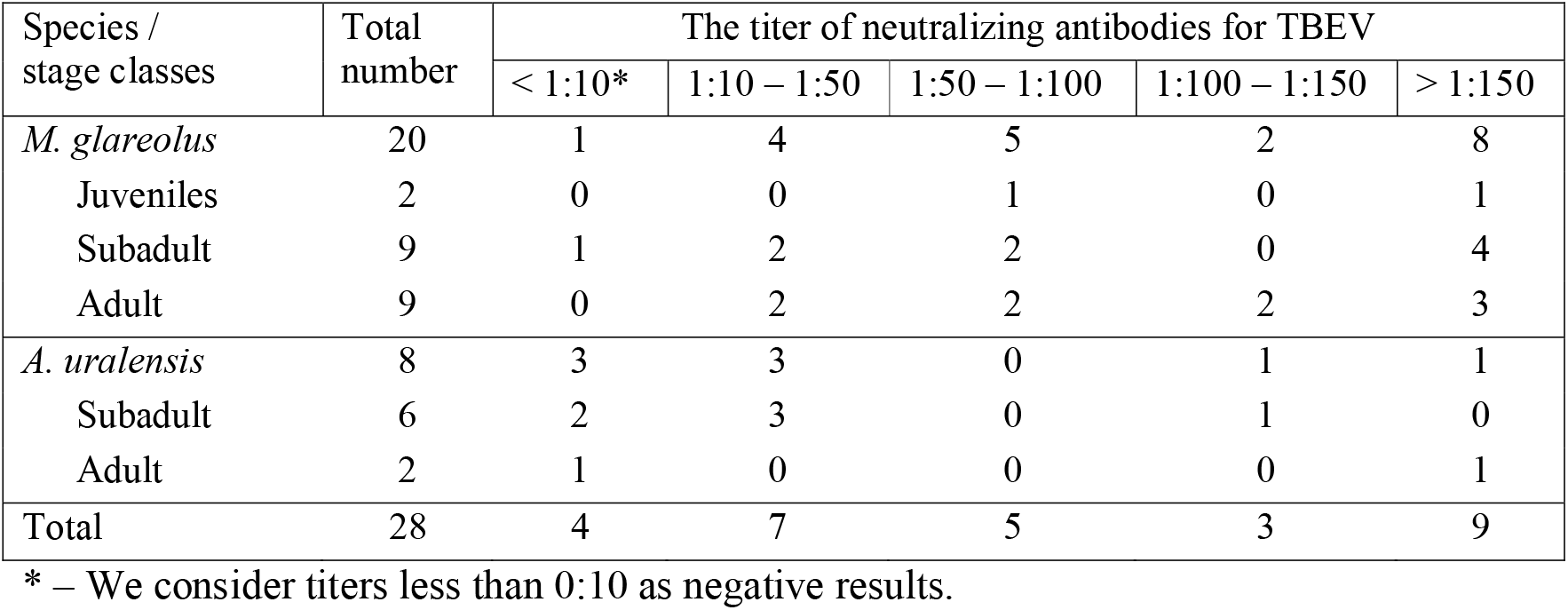
The number of animals with the corresponding titers of neutralizing antibodies for TBEV (based on a 50% plaque reduction neutralization test of the rodents’ serum)

### Virus characteristics

Six isolates of TBEV were obtained, including an isolate from the tick that bit the patient described in Case-2, four isolates from questing ticks, and one isolate from the brain tissue of a bank vole. We genetically characterized three TBEV-positive tick samples by sequencing their complete polyprotein gene. For another two tick samples, we sequenced the NS1 protein gene. The obtained isolates belong to the European subtype (Fig. 3). The sequences are most similar to the strain *Ljubljana I* (U27494), which was isolated from a patient in Slovenia in 1992 (Wallner et al., 1995). The TBEV complete polyprotein gene sequences at Moscow Forest Park had low diversity with sequence identities as high as 99.9%; the genetic distance with other strains of European subtype was approximately 2.0–2.9% (Fig. 3).

**Fig. 3.**
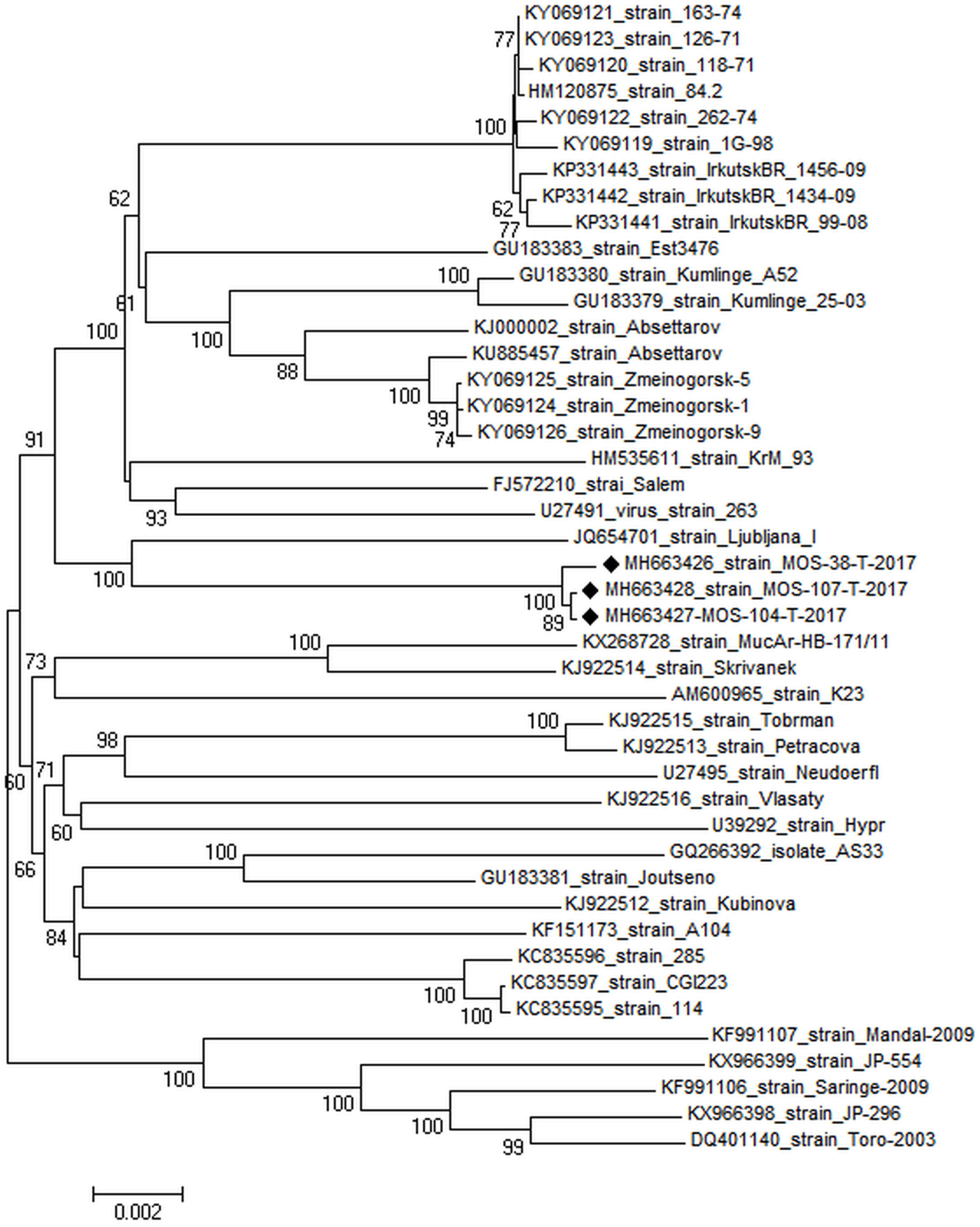
Neighbor-joining phylogenetic tree of the complete polyprotein gene (11 148 bp). The bar below indicates the number of base substitutions per site.

## Discussion

One of the important research questions is the origin of the virus in the urban forest park. As a result, we considered two hypotheses, first, the virus was introduced in the urban forest park with infected host animals (birds and/or small mammals) or with infected feeding ticks transporting by migrating birds/mammals, and secondly, the virus has been circulating here before but just had not been detected.

*I. ricinus* and *D. reticulatus* are common species for Moscow’s forest parks (Shashina and Germant, 2015; Yankovskaya et al., 2017). In Moscow Oblast the abundance of *I. ricinus* in August was up to 46,5 ticks per kilometer of flagging (Kislenko and Korotkov, 2002), the mean prevalence of *B. burgdorferi* s.l. was 19.5% (Kislenko and Korotkov, 2002; Korotkov et al., 2008). The incidence of Lyme disease in Moscow in 2015 was 9.5 cases per 100 000 residents (Yankovskaya et al., 2017). TBEV has not been detected in ticks of Moscow and Moscow Oblast (Shashina and Germant, 2015; Yankovskaya et al., 2017).

Our primary hypothesis is that the forest park was free of TBEV, and the virus was recently introduced. Consequently, three scenarios of this process are possible: first the virus was introduced with an engorged tick female on a migrating bird or small mammal. Secondly, the virus was brought into the forest park by an infected host animal with high viremia at the time of high questing activity of ticks. Alternatively, the third possibility was both of these scenarios simultaneously.

The hypothesis of recent TBEV introduction is supported by different arguments.

Previously, TBEV has not been detected in ticks or small mammals in Moscow and Moscow Oblast (“On the list of endemic territories for viral tick-borne encephalitis in 2015,” 2015). The exception is an outbreak in two districts in Northeastern part of Moscow Oblast in 1948-1957 (Drozdov, 1956). During the outbreak, about 150 cases of TBE were confirmed. Most of them were caused by consumption of raw goat or cow’s milk. At the end of the 1950s the local government had taken active measures with large-scale acaricidal spraying, which allowed to eliminate the foci. Subsequently, the TBEV was not registered in the region. It is worth mention that in 2007, one PCR-positive TBEV tick was found north of Moscow Oblast (Shevtsova et al., 2008). However, a follow-up study showed no TBEV-positive ticks in this area (Shevtsova et al., 2009).

Second, epidemiological studies of TBE in Moscow and Moscow Oblast showed that all cases were non-autochthonous, and ticks outside of Moscow Oblast bit all patients with TBE (*On the incidence of tick-borne borreliosis in Moscow in 2014*, 2015; Yankovskaya et al., 2017). Autochthonous TBE cases have not been detected since the 1950s.

Third, we found TBEV-positive ticks only at Site-1. Furthermore, mapping of the exact coordinates of the ticks’ collection sites showed the close localization of TBEV-positive ticks (Fig. 2). This evidence also supports the hypothesis of the virus’s recent introduction. If the virus had been introduced by an infected female tick, we can expect that its offspring (larvae) had been dispersed in the vicinity of the place of oviposition. The distance of the dispersion of questing nymphs and adult ticks depends on the home range size of hosts. The primary hosts for larvae of *I. ricinus* are rodents (Korenberg et al., 2013; Randolph and Craine, 1995). In the forest parks of Moscow, the most common small mammals are the bank vole (*M. glareolus*) and the pygmy field mouse (*A. uralensis*) (Tikhonova et al., 2012). We obtained the same results, and these two species also dominated in our sample. The home range of the bank vole in the breeding season varies up to 3260 m^2^ and for pygmy field mice — 1100-1800 m^2^ (Ilchenko and Zubchaninova, 1963; Kikkawa, 1964; Naumov, 1951; Zhigarev, 2004). In our study, the maximum distance between locations where the infected ticks were collected is 185 m. This is consistent with the home range size of the main hosts of immature ticks — bank voles and pygmy field mice. Therefore, the spatial distribution of the infected ticks does not contradict the hypothesis that they are the offspring of one infected female. In addition to the transovarial transmission route, TBEV can be transmitted during blood meals from a host with high viremia to feeding ticks or between cofeeding ticks (Alekseev and Chunikhin, 1990; Labuda et al., 1993). The dense occurrence of infected ticks in Site-1 does not contradict these two scenarios as well.

Fourth, only single specimens of *I. persulcatus* and *D. reticulatus* were identified in the study area. Moscow and Moscow Oblast are within the ranges of *I. ricinus, I. persulcatus* and *D. reticulatus* (Filippova, 1997, 1977). Each of these species has a mosaic distribution in the region (Korotkov et al., 2008). *I. persulcatus* in Moscow Oblast has a unimodal seasonal activity pattern with the peak in May-June that declines through August (Filippova, 1985). Therefore, we found only one female *I. persulcatus*, probably because of low activity. *D. reticulatus* has a bimodal pattern of seasonal activity, with the first peak in May and the second peak in August. We collected ticks during August-September and found only a single specimen of *D. reticulatus*. Consequently, we can conclude that this species does not form a permanent population in the forest park and that the detected specimen was accidentally found here, arriving here, apparently, with migrating birds or mammals. This evidence also highlights the hypothesis regarding the introduction of the virus.

The fifth assertion is that in the European part of Russia, three subtypes of TBEV are spread, with the Siberian subtype being predominant (Demina et al., 2009; Zlobin et al., 2003). Our results revealed that the identified virus belongs to the European subtype. The sequences of Moscow’s TBEV are most similar to the strain isolated from a patient in Slovenia in 1992 (Wallner et al., 1995). This finding can also serve as an indirect confirmation of the hypothesis that the virus was introduced from the outside; the most likely scenario is that the virus was introduced by migrating birds. The spread of TBEV with migrating birds was shown in Tomsk city (Russia), in Sweden, Norway, and Estonia (Julia Geller et al., 2013; Hasle, 2013, 2010; Moskvitina et al., 2014; Waldenström et al., 2007). The avifauna of the studied forest park includes 83 species, including the fieldfare (*Turdus pilaris*), common blackbird (*Turdus merula*), redwing (*Turdus iliacus*), song thrush (*Turdus philomelos*), thrush nightingale (*Luscinia luscinia*), European robin (*Erithacus rubecula*) and common chaffinch (*Fringilla coelebs*) (Kalyakin et al., 2014). These birds can act as tick hosts (Korenberg, 1966, 1962; Korenberg et al., 1964) and as tick vectors (Dubska et al., 2009; Hasle, 2013, 2010; Humair et al., 1993; Marsot et al., 2012; Michalik et al., 2008). The overwintering areas of the mentioned thrushes are located mainly in Italy and France (Bolshakov et al., 2009). These countries are considered low-risk areas for TBEV (European Center for Disease Prevention and Control, 2016). However, it is known that during their migration, birds use various stopover sites, where they feed and rest and where ticks may attach (Sándor et al., 2014). Therefore, birds could acquire the virus or infected ticks in Eastern Europe on the way from the overwintering grounds. Based on the arguments above, we are inclined to think that the TBEV found in this study was introduced to the forest park from other regions. Further research is needed to provide an unambiguous answer to this question.

Another important issue raised by this work is how many viral copies need to be introduced into the region for the establishment of new foci. For such tasks, the concept of *R_0_* – the basic reproduction number is useful. *R_0_* is the estimate of the average number of secondary cases produced by one primary case in a completely susceptible population (Diekmann et al., 1990). For example, in a host population of capybaras *Hydrochoerus hydrochaeris* susceptible to the tick *Amblyomma sculptum*, it was shown that the introduction of a single infected capybara or a single infected tick was unable to trigger Brazilian Spotted Fever (caused by the bacterium *Rickettsia rickettsia*) in a nonendemic area (Polo et al., 2017). Only the introduction of a single infected capybara with at least one infected attached tick was enough to trigger the disease in a nonendemic area (Polo et al., 2017). *R_0_* was defined for TBEV, but only using a model small mammal host and *I. ricinus* (Hartemink et al., 2008). The contribution of other vectors and hosts to the *R_0_* of TBEV has not been studied. In the forest park, we found not only *I. ricinus* but also *I. trianguliceps*. Ticks of *I. trianguliceps* species do not attack humans but take part in the circulation of the TBEV (Malyushina and Katin, 1965). Thus, our work highlights the need to construct more complex models for estimating *R_0_* with the inclusion of several groups of hosts (rodents and birds) and at least two vectors (*I. ricinus* and *I. trianguliceps*).

Our findings also contribute to the discussion about the role of urban areas in the development of new foci of vector-borne diseases. It is known that urban areas differ from rural areas in abiotic factors as well as in the fauna of small mammals and birds and their respective behaviors (home-range size, wintering, migration, etc.) (Forman, 2014; Klausnitzer, 1993). Therefore, the role of urban areas in the emergence of new foci requires special attention.

## Acknowledgments

This research did not receive any specific grant from funding agencies in the public, commercial, or not-for-profit sectors.

We thank Novik, I.V. and Midyannik, G.A. for assistance with ticks collection, and Uspensky, I.V. for comments that greatly improved the manuscript. We would also like to thank Antonovskaya, A.A. (Lomonosov Moscow State University) for her help in identifying of chiggers species.

## Notes

#### Summary of Updates

We corrected an error in the mention of the species of wood mice: instead of Apodemus sylvaticus, we indicated Apodemus uralensis.

